# Effects of food availability cycles on phase and period of activity/rest rhythm in *Drosophila melanogaster*

**DOI:** 10.1101/2021.11.17.468936

**Authors:** Viveka Jagdish Singh, Sheetal Potdar, Vasu Sheeba

**Author notes:** Molecular Neurobiology Laboratory, Salk Institute for biological studies, La Jolla, CA 92037. Corresponding author: Sheeba Vasu **Address**: Chronobiology and Behavioural Neurogenetics Laboratory, Neuroscience Unit2, Jawaharlal Nehru Centre for Advanced Scientific Research, Jakkur P.O., Bangalore – 560 064 **Email**.

## Abstract

Foraging and feeding are indispensable for survival and their timing depends not only on the metabolic state of the animal but also on the availability of food resources in their environment. Since both these aspects are subject to change over time, these behaviours exhibit rhythmicity in occurrence. As the locomotor activity of an organism is related to its disposition to acquire food, and peak feeding in fruit flies has been shown to occur at a particular time of the day, we asked if cyclic food availability can entrain their rhythmic activity. By subjecting flies to cyclic food availability *i.e*., feeding/starvation (FS) cycles, we provided food cues contrasting to the preferred activity times and observed if this imposed cycling in food availability could entrain the activity/rest rhythm. We found that phase control, which is a property integral to entrainment, was not achieved despite increasing starvation duration of FS cycles (FS12:12, FS10:14 and FS8:16). We also found that flies subjected to T21 and T26 FS cycles were unable to match period of the activity rhythm to short or long T-cycles. Taken together these results show that external food availability cycles do not entrain the activity/rest rhythm of fruit flies. However, we find that starvation induced hyperactivity causes masking which results in phase changes. Additionally, T-cycle experiments resulted in minor period changes during FS treatment. These findings highlight that food cyclicity by itself may not be a potent zeitgeber but may act in unison with other abiotic factors like light and temperature to help flies time their activity appropriately.

## INTRODUCTION

Circadian clocks integrate cues from the environment and temporally regulate physiological and behavioural programs to aid animals fulfill their daily needs by anticipating cyclic changes in their day-to-day environment and time their physiology accordingly (Dunlap, 2003). This is achieved by the process of entrainment which is the ability of the clock to synchronize to cyclic cues in the environment. Abiotic factors that cycle with time of day such as light and temperature serve as time cues or “zeitgebers” to the clock (Pittendrigh, 1960); (Johnson et al., 2003). Similarly, biotic factors such as food resources may also serve as time cues to the clock of many animals (Hau and Gwinner, 1996); (Sharma et al., 2000); (Frisch and Aschoff, 1987).

Food resources frequently undergo changes in quality and quantity. While these changes are more apparent over seasons, daily food availability may also change as is documented in the case of various plant-pollinator and prey-predator interactions (Bloch et al., 2017); (Kronfeld-Schor et al., 2017). The cyclicity of these interactions are made possible by the circadian clocks that regulate diverse aspects of pollination such as those involved in flower advertisements in plants (Overland, 1960); (Matile, 2006); (Fenske et al., 2015); (Yon et al., 2016) and foraging activities in pollinator species (Fenske et al., 2018). Activity of several insect species such as bees (Bloch et al., 2017), mosquitoes (reviewed in (Sougoufara et al., 2017)), bedbugs (Romero et al., 2010) (reviewed in (Barrozo et al., 2004)) have been shown to be influenced by food availability.

Restricted food access in rodent models under laboratory conditions invokes an anticipatory response in the form of an activity bout before food availability called the food anticipatory activity (FAA) (Richter, 1922). FAA occurs for as long as food is restricted and even at times when animals are not usually active. For example, when food is restricted to daytime, FAA is observed during daytime which is otherwise a period of low activity in nocturnal animals (Mistlberger, 2011); (Carneiro and Araujo, 2012). FAA is not dependent on the canonical light entrainable oscillator (LEO) located in the Suprachiasmatic Nucleus (SCN) (Stephan et al., 1979), and is thought to be controlled by another clock which is termed Food entrainable oscillator (FEO) which has not yet been localized (Pendergast and Yamazaki, 2018). Furthermore, it has been shown that feeding entrains a peripheral clock in the liver (Damiola et al., 2000). Therefore, food provided at unusual times of the day can disrupt the phase relationship between the LEO and the liver peripheral clock. Other than the occurrence of FAA, the overall activity/rest pattern of these animals remains largely unaffected by change in food availability when SCN is intact. However, in SCN-lesioned animals, FEO can completely entrain activity rhythms to food availability cycles (Pendergast and Yamazaki, 2018), suggesting that food can act as a secondary zeitgeber

In the fruit fly, *Drosophila melanogaster* a peripheral clock in olfactory receptor neurons of the antenna regulates a circadian rhythm in olfactory responses with a peak in the middle of the night (Krishnan et al., 1999); (Tanoue et al., 2004). Similarly, a diurnal rhythm in electrophysiological responses of the labellar gustatory receptor neurons (GRN) has been reported with a peak in the morning hours. The GRN clock also regulates a behavioural gustatory rhythm in proboscis extension reflex (PrER); an appetitive behaviour with a peak in the morning (Chatterjee et al., 2010). Fruit flies also feed rhythmically with a peak in the morning in light/dark cycles and early subjective day in constant conditions (Xu et al., 2008). All these rhythms in *Drosophila* have been shown to be controlled by peripheral clocks while the activity/rest rhythm is known to be regulated by the central clock neurons (Helfrich-Forster, 1998). However, whether and how food availability affects any of these rhythms is unknown.

An organism’s active phase is the time when most of the resource gathering and energy requirements are likely to be fulfilled. Hence locomotion of most animals is a function of various drives such as foraging, feeding, mating, oviposition etc. It is imperative to bring congruence between the internal drive to feed and the availability of food resources in the environment. Circadian clocks could facilitate this by adjusting the active phase of the organism such that the animal performs foraging and feeding behaviours while food is likely to be available in the environment. Since being in an active state is closely tied to an organism’s disposition to acquire food resources, we asked if changing the time of food availability can bring about changes in the activity patterns and affect the underlying clock in *D. melanogaster*. In this study, we test the hypothesis that food availability cycles can act as a zeitgeber in entraining the activity/rest rhythm of *Drosophila melanogaster* by imposing various types of feeding cycles.

## METHODS

### Locomotor Activity Assay

Locomotor activity rhythm of flies was recorded using the *Drosophila* Activity Monitor (DAM, Trikinetics, USA). 4–5-day old virgin male flies, unless mentioned otherwise were recorded in LD 12:12 with *ad libitum* food (standard cornmeal medium) at 25°C for 2-3 days following which Feeding: Starvation (FS) cycles were imposed in constant dark (DD). The period of starvation lasted for either 12, 14 or 16 h depending upon the regime. Experimental flies received standard cornmeal food during the ‘feeding’ phase of the cycle and were transferred into 2% agar during the ‘starvation’ phase. FS cycles were imposed for 7 days following which flies were shifted to DD *ad libitum* food (DD *ad lib*) conditions for the next 7 days. Age matched flies that were transferred into fresh food tubes at the same time as experimental flies served as disturbance controls. All transfers were conducted under far-red light illumination (>630nm) in DD.

For the phase-shifted FS cycle experiment, 5–6-day old flies were subjected to the first FS (FS1) for 5 days after which the second FS (FS2) was imposed by either advancing or delaying the food transfers by 6 h with respect to FS1 for a period of 7 days. Acrophase (calculated using Actogram J, (Schmid et al., 2011)) was used as a phase marker to obtain the phase of the rhythm during and after FS cycles in all the regimes.

For T26 and T21 cycles, 5–6-day old virgin male flies were recorded in LD 12:12 with *ad libitum* food (standard cornmeal medium) at 25°C for 5 days following which FS cycles were imposed in constant dark. A T26 FS cycle was imposed on these flies such that the flies experienced 13 h of food availability and 13 h of starvation. Similarly, a T21 FS cycle was imposed where flies experienced 10.5 h of food availability and 10.5 h of starvation. Age matched flies served as disturbance controls as previously described. Seven such cycles were imposed following which flies were shifted to DD *ad libitum* food conditions for the next 7 days. A *chi-square* periodogram analysis was done (using ClockLab software, Actimetrics, Wilmette, IL, USA) to determine periodicity during T26 and T21 feeding regimes as well as during DD *ad libitum* phase. In all the assays, flies were reared under LD12:12 regime before the start of the assay and all experiments were conducted at 25°C.

### Statistical analyses

Daily acrophases were compared using repeated measures ANOVA with day as the repeated measure and treatment as the between-group factor. Mauchly’s test for sphericity was performed on all the data sets and Greenhouse-Geisser corrections were applied when the assumption for sphericity was not met. The above tests were performed using IBM, SPSS Statistics for windows (version 26, 2019, IBM corp., Armonk, N.Y., USA). Multiple *post hoc* pairwise comparisons were performed using *t-*tests with Bonferroni corrections. Activity levels during starvation for all the FS regimes were analysed similarly on IBM, SPSS. Inter-individual phase synchrony between controls and experimental flies was tested by measuring the degree of dispersion of mean phases averaged across last 3 days of the FS cycles. Wallraff Rank sum test for angular dispersion was performed on phase values (radians) using R core team (version 3.6.1, 2019, R: A language and environment for statistical computing. R Foundation for Statistical Computing, Vienna, Austria. URL http://www.R-project.org/). Δphase – defined as change in phase on the first day of DD *ad lib* from the mean acrophase (last 3 days) during FS regime, was compared using one sample *t*-test against a reference constant 0. Additionally, two sample *t*-test was used to compare Δ phases between controls and experimental flies. For T21 & T26 experiments, a Chi Square test for proportions was performed using GraphPad Prism (version 9.2.0 for Windows, GraphPad software, San Diego, California, USA, www.graphpad.com) to compare proportions of flies exhibiting different periodicities. Change in period was tested using Mann-Whitney U test. All other analyses were performed on STATISTICA (version 7, 2004, StatSoft Inc, Tulsa, OK, USA).

## RESULTS

In order to address if external food availability cycles or the Feeding: Starvation (FS) cycles entrain the activity/rest rhythm of *D. melanogaster*, we subjected flies to three different FS cycles of increasing starvation duration – namely, FS12:12 (12 h of feeding followed by 12 h of starvation, Fig 1A), FS10:14 (10 h of feeding followed by 14 h of starvation, Fig 1B) and FS8:16 (8 h of feeding followed by 16 h of starvation, Fig 1C). Activity/rest rhythms were recorded in LD12:12 at 25°C on *ad libitum* food for 2-3 days before shifting to one of the aforementioned FS regimes in constant dark (DD). In all three FS regimes, the feeding duration overlapped, partly or completely with daytime in the previous LD cycles. Following 7-8 days of FS regime, flies were subjected to DD with *ad libitum* food (DD *ad lib*). Disturbance control and experimental flies show startle bouts of activity when they are moved to new tubes (Fig 1A-C, left, arrows). While these startle bouts can be attributed to disturbance due to change of tubes, we expected to observe changes in the activity/rest rhythm because of FS cycles over and above the disturbance caused during the assay (Fig 1). For example, as expected from previous studies (Connolly, 1966), experimental flies show increased activity level during starvation compared to control flies (Fig 1A-C, right). To determine if FS cycles are indeed entraining the activity/rest rhythm we examined classical criteria of entrainment, namely – day-to-day phase stability, inter-individual phase synchrony, phase control and period matching with zeitgeber cycle (τ = T) (Moore-Ede et al., 1982). We used acrophase which is the radial centre of mass of activity (Diez-Noguera, 2013) as the phase marker in all the experiments.

**Figure 1:**
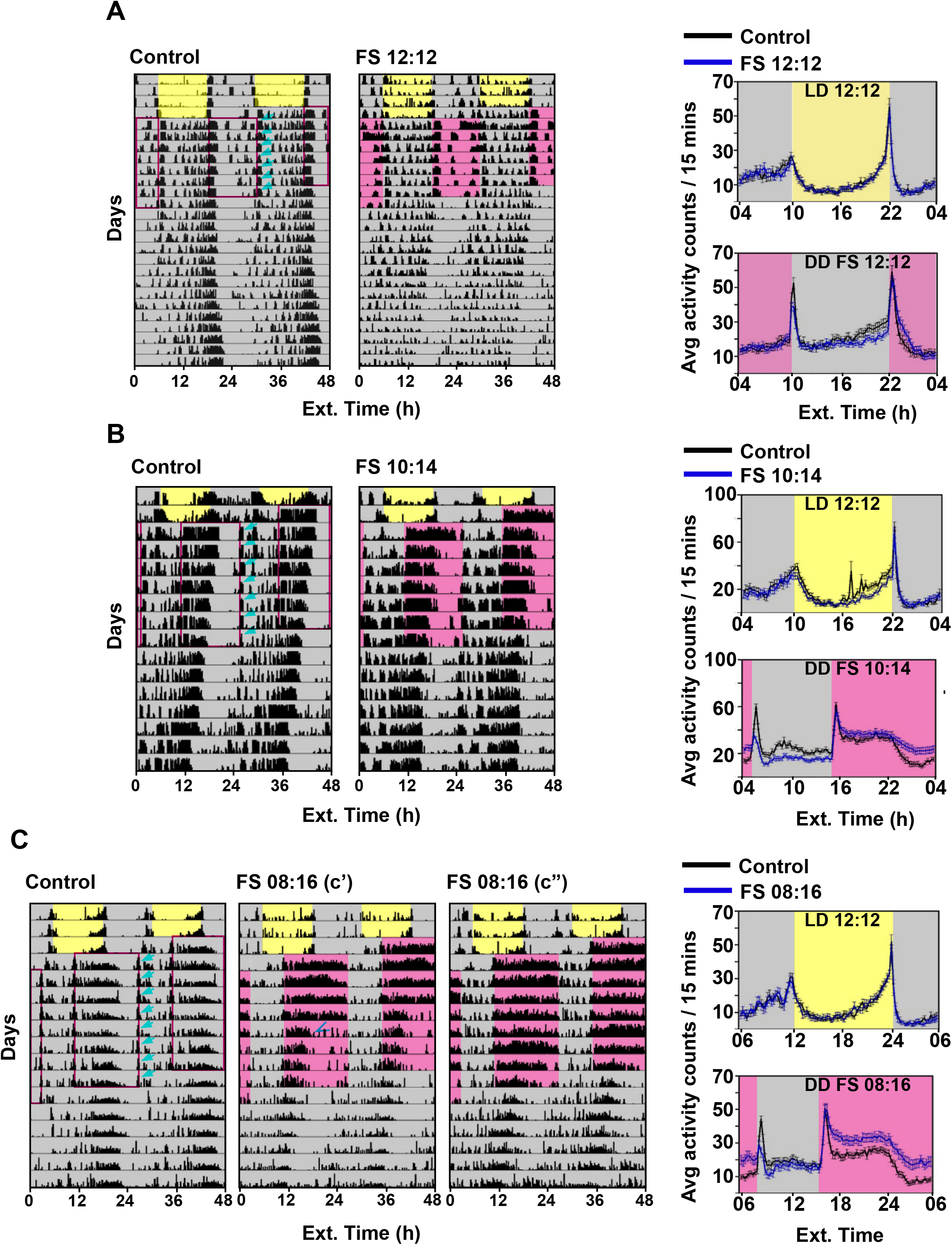
Flies subjected to FS cycles display a combination of free running and masking behaviour. Representative double plotted actograms of **(left)** age matched disturbance controls and flies subjected to FS cycles; **(A)** FS12:12, **(B)** FS10:14, **(C)** FS8:16 and **(right)** average profiles of activity binned in 15 min intervals **(top)** before and **(bottom)** during FS cycles, **(A)** FS12:12, **(B)** FS10:14, **(C)** FS8:16. Error bars for activity profiles are ± standard error of the mean (SEM). Yellow shaded region represents daytime and grey shaded region represents night-time with *ad libitum* food. Pink shaded region represents starvation. In every regime, disturbance caused due to transfer of flies in fresh tubes results in small bursts in activity, arrows indicate the startle bouts. In FS 8:16, (c’) a small fraction of flies (7/23) shows gradual reduction in activity (arrowhead) during the starvation window across days of the treatment; (c’’) another fraction of flies (14/23) displays elevated activity levels across all days of the treatment. *x*-axis indicates external time.

### Day-to-day phases vary in flies subjected to FS cycles of 10:14 and 8:16

We tested day-to-day stability of phases to determine stable phase relationship with zeitgeber cycles. To compare daily phases, we performed repeated measures ANOVA on acrophases of control and experimental flies during FS treatment with day as the repeated measure and treatment as a between-group fixed factor. We found that daily acrophases of flies subjected to FS12:12 were similar to those of control flies on all cycles. Further, acrophases of both groups on first two cycles were different from acrophases in subsequent cycles suggesting phase changes due to startle responses in both control and experimental flies (Fig 2A, repeated measures ANOVA, Greenhouse-Geisser ε = 0.56, F_(3.34, 176.92)_ = 8.93, main effect of day, *p* < 0.001 followed by pair-wise *t*-tests with Bonferroni corrections for 21 comparisons). In FS10:14 regime, on day 1, controls show a significantly different phase compared to phases on subsequent cycles 2 and 3 suggesting that day 1 phase is affected by disturbance (Fig 2B, Greenhouse-Geisser ε = 0.66, F_(3.98,222.99)_ = 5.66, day × treatment, *p* < 0.001, followed by pair-wise *t*-tests with Bonferroni corrections for 49 comparisons). However, it stabilizes within a day as phases from the 2nd to the 7th cycle remain unchanged. Experimental flies under FS10:14 show gradual changes in day-to-day phases. Acrophases in the first two cycles of the FS regime are significantly different from the last two cycles of the treatment (Fig 2B). Furthermore, we also observed that experimental flies have acrophases which are significantly different from controls in the first half of the treatment (cycles 1-3) whereas these acrophases start resembling acrophases of controls in the second half of the treatment (cycles 4-7). This suggests that imposing FS10:14 cycles result in transient phase changes which gradually disappear after a few cycles.

**Figure 2:**
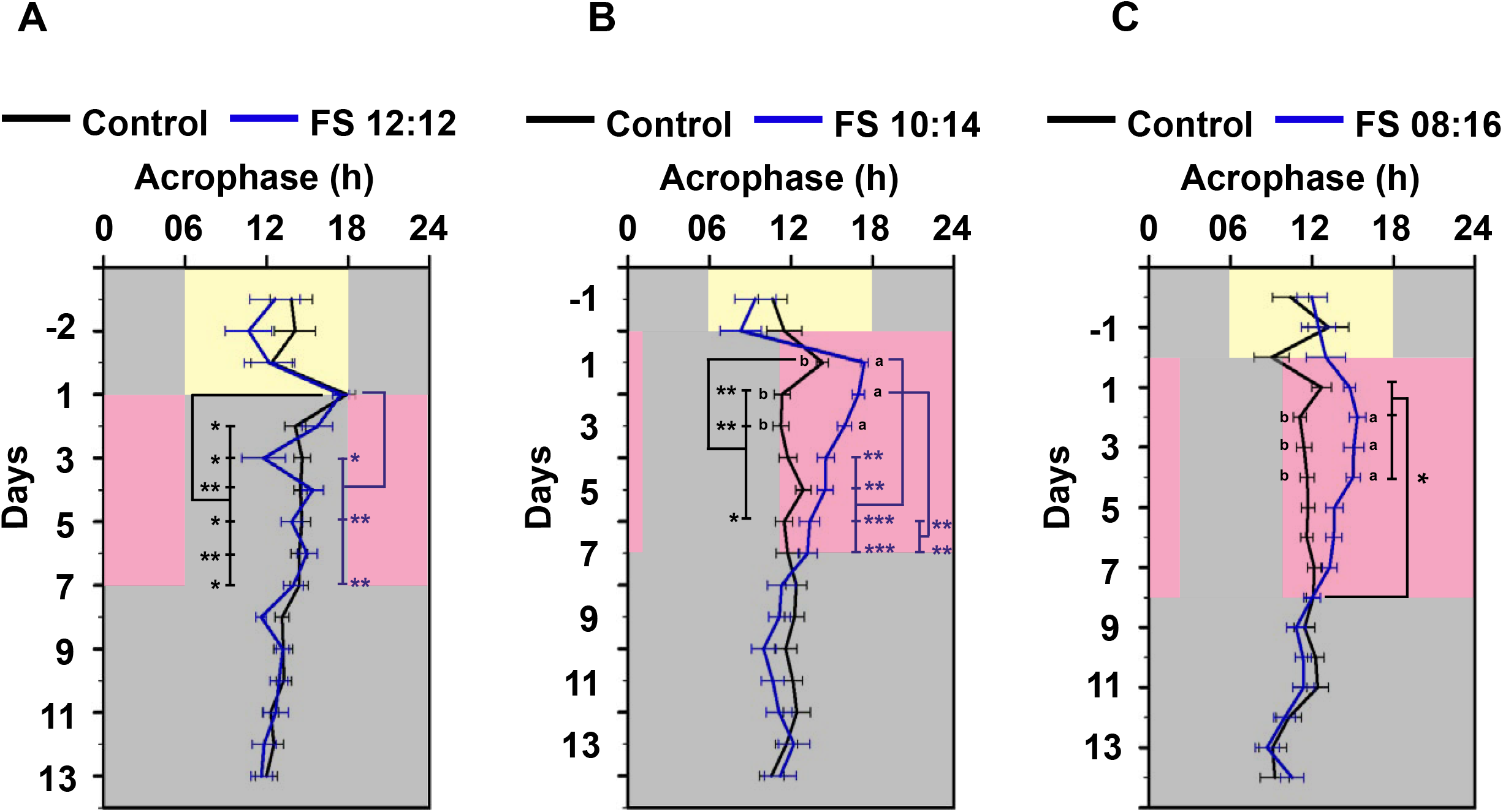
Acrophases change across days in flies experiencing FS10:14 & FS8:16. Mean acrophases ± SEM of controls (black line) and FS flies (blue line) across days under LD 12:12, FS treatment and DD *ad lib*, **(A)** FS12:12, **(B)** FS10:14 and **(C)** FS 8:16. Error bars are standard error of the mean. Experimental flies and their controls show phase change on day 1 of the regime. Controls revert to previous phase immediately while experimental flies gradually return to phases similar to controls. Letters denote significant differences (p < 0.05) between controls and FS flies. *, **, *** denote significant differences across days with *p* < 0.05, 0.01, 0.001 respectively.

Similarly, phases of the experimental flies experiencing FS8:16, also change gradually across days with acrophases on the first and second cycles being significantly different from acrophase on last day of the treatment (8th cycle, Fig 2C, Greenhouse-Geisser ε = 0.74, F_(5.16,237.45)_ = 5.26, day × treatment, *p* < 0.001, followed by pair-wise *t*-tests with Bonferroni corrections for 64 comparisons). On the other hand, control flies do not show any phase changes across all cycles. Additionally, similar to FS10:14, acrophases of FS8:16 flies are significantly different from controls from the beginning of the treatment (cycles 2-4) but start resembling acrophases of the controls towards the end of the treatment (cycles 5-8).

Controls in all the regimes are affected by disturbance on the first day of FS cycle after which they show phase stability. On the other hand, phases of experimental flies are affected on the first few FS cycles which take longer to attain phases similar to the controls. Given that the internal period is not too different from 24 h (Table 1), it is difficult to ascertain whether this day-to-day phase stability in the latter half of FS treatment is due to free-running, masking or entrainment. Hence, we used other measures to differentiate between these possibilities.

**Table:1.** Period and amplitude values of control and experimental flies during and after FS 12:12, FS 10:14 and FS 8:16 cycles.

| **MALES** |  |  |  |  |  |  |  |  |
| --- | --- | --- | --- | --- | --- | --- | --- | --- |
| Regime | Period |  | Power |  | n |  | % Rhythmicity |  |
|  | Control | Expt. | Control | Expt. | Control | Expt. | Control | Expt. |
| During FS 12:12 1 | 23.8 ± 0.05 | 23.7 ± 0.05 | 167.07 ± 6.9 | 166.48 ± 5.7 | 29 | 32 | 100 | 100 |
| During FS 12:12 2 (phase delayed) | 24.02 ± 0.04 | 23.99 ± 0.02 | 203.7 ± 8.4 | 190.04 ± 7.2 | 29 | 32 | 100 | 100 |
| DD *ad lib* after FS 12:12 | 23.83 ± 0.07 | 23.72 ± 0.07 | 170.8 ± 12.6 | 156.5 ± 12.2 | 28 | 30 | 96.43 | 96.67 |

| Regime | Period |  | Power |  | n |  | % Rhythmicity |  |
| --- | --- | --- | --- | --- | --- | --- | --- | --- |
|  | Control | Expt. | Control | Expt. | Control | Expt. | Control | Expt. |
| During FS 10:14 | 23.91 ± 0.04 | 23.84 ± 0.06 | 205.3 ± 6.6 | 229.7 ± 8.8 | 30 | 28 | 100 | 100 |
| DD *ad lib* after FS 10:14 | 23.67 ± 0.04 | 23.62 ± 0.08 | 210.38 ± 14.4 | 237.94 ± 16.1 | 30 | 27 | 87.09 | 88.89 |
|  | Control | Expt. | Control | Expt. | Control | Expt. | Control | Expt. |
| During FS 08:16 | 23.97 ± 0.02 | 23.95 ± 0.02 | 249.9 ± 9.43 | 243.85 ± 10.1 | 32 | 23 | 100 | 100 |
| DD *ad lib* after FS 08:16 | 23.76 ± 0.09 | 23.70 ± 0.08 | 120.11 ± 6.6 | 122.31 ± 6.6 | 32 | 23 | 87.5 | 95.65 |

| **FEMALES** |  |  |  |  |  |  |  |  |
| --- | --- | --- | --- | --- | --- | --- | --- | --- |
| Regime | Period |  | Power |  | n |  | % Rhythmicity |  |
|  | Control | Expt. | Control | Expt. | Control | Expt. | Control | Expt. |
| During FS 12:12 1 | 23.8 ± 0.07 | 23.99 ± 0.14 | 131.04 ± 5.28 | 126.49 ± 4.9 | 28 | 29 | 85.71 | 93.1 |
| During FS 12:12 2 (phase delayed) | 23.89 ± 0.26 | 24.46 ± 0.41 | 155.65 ± 10.52 | 140.87 ± 7.19 | 26 | 27 | 73.08 | 81.5 |
| DD *ad lib* after FS 12:12 | 23.96 ± 0.11 | 23.6 ± 0.16 | 125.75 ± 11.3 | 103.5 ± 8.52 | 23 | 25 | 52.2 | 26.09* |

| Regime | Period |  | Power |  | n |  | % Rhythmicity |  |
| --- | --- | --- | --- | --- | --- | --- | --- | --- |
|  | Control | Expt. | Control | Expt. | Control | Expt. | Control | Expt. |
| During FS 10:14 | 24 ± 0.02 | 23.92 ± 0.05 | 164.8 ± 7.3 | 179.2 ± 9.8 | 25 | 25 | 100 | 100 |
| DD *ad lib* after FS 10:14 | 23.6 ± 0.15 | 23.4 ± 0.54 | 169.18 ± 13.23 | 152.8 ± 10.93 | 20 | 25 | 60 | 40 |
|  | Control | Expt. | Control | Expt. | Control | Expt. | Control | Expt. |
| During FS 08:16 | 24.15 ± 0.1 | 24.3 ± 0.24 | 160.5 ± 9.07 | 215.77 ± 13.1 | 24 | 25 | 95.83 | 96 |
| DD *ad lib* after FS 08:16 | 23.75 ± 0.21 | 23.36 ± 0.18 | 98.4 ± 6.6 | 98.65 ± 9.53 | 20 | 16 | 40 | 37.5 |

### Feeding: Starvation cycles do not increase inter-individual phase synchrony

An entrained rhythm is phase-locked to the zeitgeber cycle, and this phase relationship is stable across multiple cycles and reproducible across individuals. This implies that in an entrained condition, individual flies will exhibit similar phases resulting in higher inter-individual synchrony. Here we examined the extent of phase dispersion within control and experimental fly groups during FS regimes. If FS cycles were entraining the activity/rest rhythm we would expect a smaller dispersion with greater consolidation of phases under entrained conditions. We find that the degree of dispersion of acrophases of flies in each of the experimental regimes of FS12:12 (Fig 3A, Wallraff rank sum test for angular dispersion, Kruskal-Wallis Chi Square = 2.32, *df* = 1, *p* = 0.13), FS10:14 (Fig 3C, Kruskal-Wallis Chi Square = 0.007, *df* = 1*, p* = 0.93) and FS8:16 (Fig 3E, Kruskal-Wallis Chi Square = 1.31, *df* = 1, *p* = 0.25) was not statistically different from their respective disturbance controls suggesting that FS cycles are not efficient in synchronizing the phases of experimental flies.

**Figure 3:**
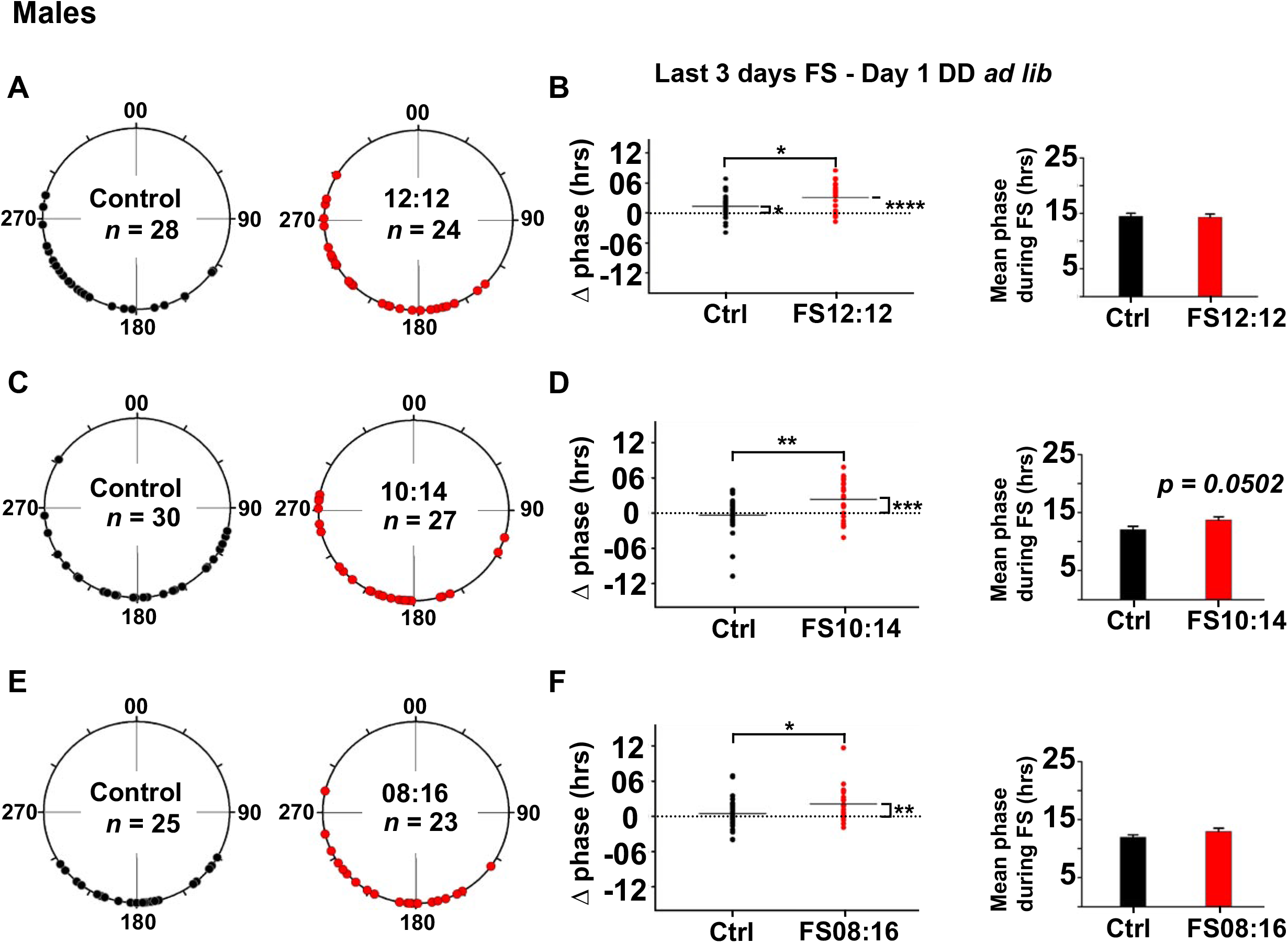
FS cycles are inefficient in exerting phase control and do not result in greater phase synchrony among flies. **(A)** Polar plots depicting average acrophases of individuals for the last 3 days duringFS12:12. black circles indicate phases of control flies and red circles indicate phases of experimental flies. Zero degrees in each polar plot was set to ZT18 of the previous LD cycles. **(B-Left)** Scatter plots depicting showing differences in acrophases (Δ phase) of individual control (black) and experimental (red) flies; horizontal line represents the mean Δ phase. Horizontal dotted lines at 0 indicates no change in phase. **(B-Right)** Bar graph depicts mean acrophase ± SEM (h) of disturbance controls (black) and experimental (red) flies on the last 3 days. Degree of phase dispersion, phase change and mean phase in **(C, D)** FS10:14 and (E, F) FS8:16 respectively, all other details same as **(A & B)**. *,** indicate *p* < 0.05, 0.01 respectively.

Additionally, we asked whether abrupt shifts in food availability schedules may reveal any features that were not previously detectable. Two jetlag FS12:12 experiments were conducted, where the second set of FS cycles (FS2) were either phase delayed (Supp Fig 1A) or advanced (Supp Fig 1B) with respect to previous FS cycle (FS1). Similar to the previous FS12:12 (Fig 1), the activity rhythm of the experimental flies during FS1 and FS2 was similarly phased as their disturbance controls (Supp Fig 1). The degree of phase dispersion during the first FS cycle and second FS cycle was also not different in experimental flies as compared to their controls in both phase delay (Fig 4A, FS1-Kruskal-Wallis Chi Square = 0.9, *df* = 1, *p* = 0.77, FS2-Kruskal-Wallis Chi Square = 1.67, *df* = 1, *p* = 0.2) and advance conditions (Fig 4B, FS1-Kruskal-Wallis Chi Square = 0.88, *df* = 1, *p* = 0.35, FS2-Kruskal-Wallis Chi Square = 1.33, *df* = 1, *p* = 0.25). These results suggest that FS cycles fail to increase inter-individual synchrony of activity rhythms.

**Figure 4:**
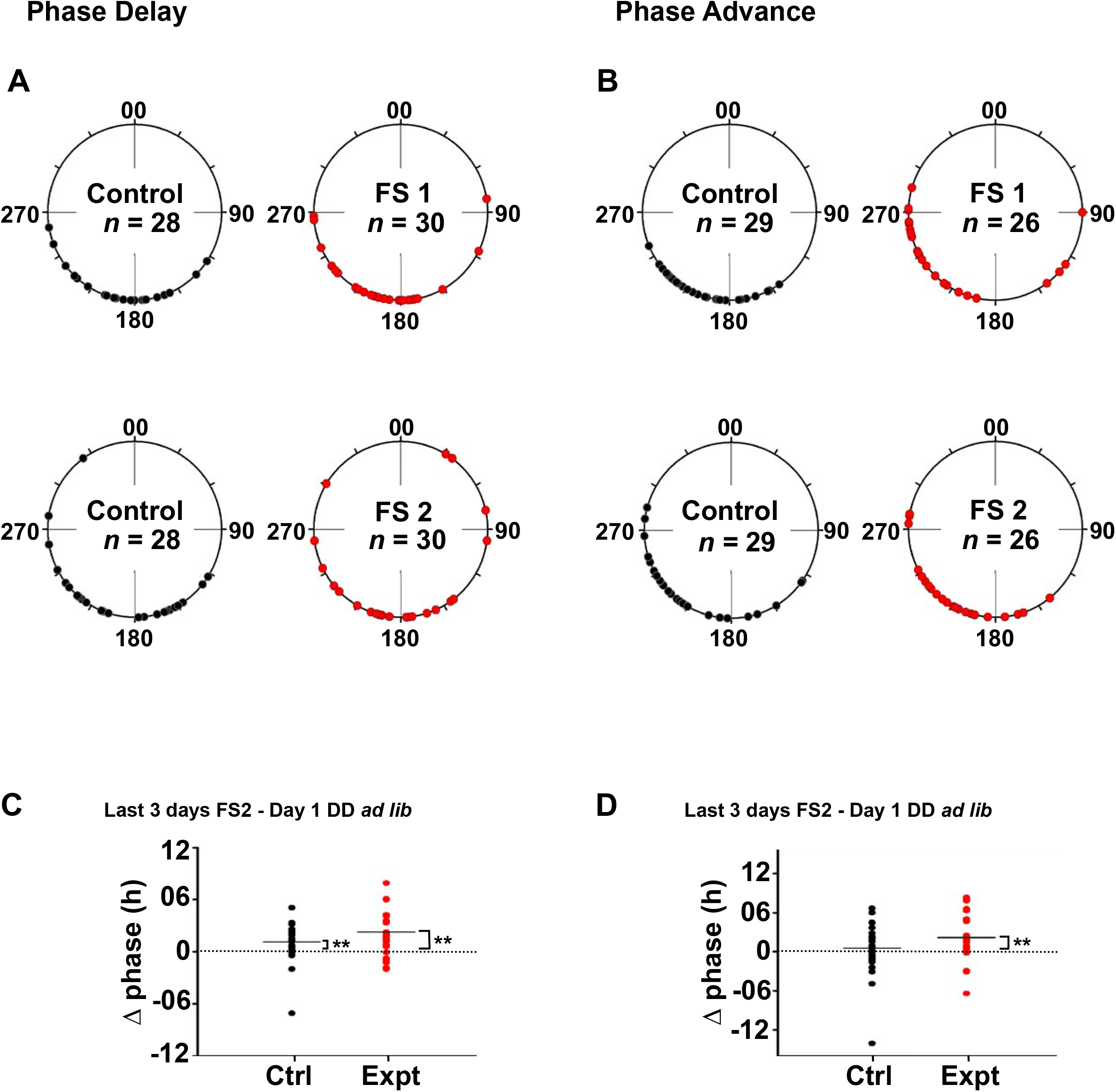
Phase shifted FS cycles do not reveal any difference in inter-individual phase synchrony. **(A)** Polar plots depicting averaged phases for the last 3 days of individual flies during FS12:12. (**Left)** black circles indicate phases of control flies and **(right)** red circles indicate the phase of flies subjected to the first FS12:12 (FS1). **(Bottom)** Polar plots depicting average phases (last 3 days) of **(left)** control and **(right)** experimental individuals during FS2, which was phase ***delayed*** by 6 hours compared to FS1. The degree of dispersion of acrophases in disturbance controls and experimental flies was not significantly different during FS1 and FS2. **(B)** Polar plots depicting averaged phases for the last 3 days of individual flies during the first FS12:12 regime (FS1). Polar plots depicting average phases (last 3 days) of control and experimental individuals during FS2, which was phase ***advanced*** by 6 hours compared to FS1. The degree of dispersion of acrophases in disturbance controls and experimental flies was not significantly different during FS1 and FS2. Scatter plots showing Δ phase for individual control (black) and experimental (red) flies when phase on the first day in DD *ad libitum* was subtracted from averaged phase in the last 3 days of **(C)** phase delayed and **(D)** advanced FS2 cycles *,** indicate *p* < 0.05, 0.01 respectively. Other details as in Figure 3.

### Feeding: Starvation cycles of different starvation durations do not exert phase control on activity/rest rhythm

To test if the phase in constant conditions follows from the previously entrained phase (phase control), the change in acrophase (Δ phase) was quantified by subtracting acrophase on the first day of DD *ad lib* from the mean acrophase (last 3 days) during FS regime. In case of phase control, the expectation is that Δ phase will not be significantly different from zero. We found that the Δ phase was significantly different from zero in both the controls as well as FS12:12 flies suggesting that the disturbance itself brings about some change in phase in both the groups (Fig 3B, left, experimental t_(27)_ = 5.73, *p* < 0.001, One sample *t-*test against reference constant 0 and control t_(26)_ = 2.76, *p* = 0.01). However, Δ phase in the experimental flies is significantly higher than the controls implying that the FS cycles fail to exert phase control (Fig 3B, left t_(53)_ = 2.45, *p* = 0.018, Two sample *t-*test). Additionally, when two FS12:12 cycles were imposed consecutively (FS1 & FS2), either phase delayed (Fig 4C, Supp Fig 1A) or phase advanced (Fig 4D, Supp Fig 1B) with respect to one another, the phase of the activity/rest rhythm continued to follow from the previous entrained phase in LD12:12 irrespective of FS2 suggesting lack of phase control (Fig 4C, D, Supp Fig 1) even in these regimes. Here again, Δ phase was significantly different from zero in experimental flies experiencing phase shifted FS cycles suggesting lack of phase control (Fig 4C, t_(29)_ = 3.37, *p* = 0.002, One sample *t*-test against reference constant 0 and Fig 4D, t_(27)_ = 3.29, *p* = 0.0028, One sample *t*-test against reference constant 0) as well as controls in the phase delay regime suggesting some disturbance related effects (Fig 4C, t_(27)_ = 2.88, *p* = 0.007, One sample *t*-test against reference constant 0). Even though control flies of FS12:12 show Δ phase value which is significantly different from zero, we found that the PERIOD (PER) accumulation rhythm in the ventrolateral neurons (LNv) neurons of individual fly brains does not undergo any change because of disturbance alone (*data not shown*) suggesting that disturbance does not affect the central clock.

When the duration of starvation was increased to 14 h in FS10:14, or to 16 h in FS8:16, Δ phase was significantly different from zero in the experimental flies but not in controls (Fig 3D, left, t_(28)_ = 4.14, *p* = 0.0002, One sample *t-*test against reference constant 0 and Fig 3F, left t_(22)_ = 3.6, *p* = 0.0015, One sample *t-*test against reference constant 0). This suggests that lack of phase control persists in experimental flies despite increasing starvation. In addition, Δ phase of experimental flies experiencing FS10:14 and FS8:16 was significantly higher than the controls (Fig 3D, left, t_(56)_ = 3.34, *p* = 0.0015, Two sample *t-*test and Fig 3F, left, t_(46)_ = 2.035, *p* = 0.0475, Two sample *t-*test). In FS10:14 the mean phase during the treatment is marginally different between the control and experimental flies (Fig 3D, right, t_(56)_ = 2.0007, *p* = 0.0502, Two sample *t-*test). The change in mean phase during FS can be attributed to acrophases affected by <u>s</u>tarvation induced <u>h</u>yperactivity (SIH, Fig 1B, right).

In all the FS regimes tested, we found that Δ phase was different from zero in all experimental flies (and also in FS12:12 controls). This suggests that the stable phase attained during the last three FS cycles changed quickly on the first day of DD *ad lib*, which is characteristic of masking. SIH was observed on all days among experimental flies which may influence acrophases resulting in masking (Fig 1, right, Supp Fig 3). Previously we had seen that FS regime induces some phase changes that gradually return to phase values similar to controls during the latter half of the regime which may reflect free running phases in flies experiencing FS. This suggests that phases during the FS cycles were possibly intermediary between the internal clock and masking components. Overall, lack of phase control and lower inter-individual synchrony suggests that FS cycles may not bring about stable entrainment in activity/rest rhythms in *Drosophila melanogaster*.

Since female flies feed more compared to males (Wong et al., 2009), we asked if FS cycles can entrain activity/rest rhythm in female flies. We found similar results when female flies were subjected to FS8:16 wherein most flies showed excessive activity in the starvation window (Supp Fig 2A). Average phase was significantly different from the controls during FS8:16 which immediately reverted to pre-FS phase after the FS treatment (DD *ad lib*) (Supp Fig 2B, repeated measures ANOVA with day as the repeated measure and treatment as between-group fixed factor was performed (main effect of treatment F_(1, 47)_ = 27.51, *p* < 0.001); C-centre t_(47)_ = 2.66, *p* = 0.0105, Two sample *t*-test). Additionally, we found that there was no phase control -as Δ phase was significantly different from zero (Supp Fig 2C, left, t_(24)_ = 4.96, *p* < 0.001, One sample *t*-test against reference constant 0) nor was there inter-individual synchrony (Supp Fig 2C-right, Wallraff rank sum test for angular dispersion). Altogether, these results indicate that activity rhythms of both males and females do not entrain to FS cycles.

### Starvation Induced hyperactivity is not sustained across FS cycles

Starvation has been shown to result in excess activity levels which fall to the baseline levels after food has been provided (Connolly, 1966); (Yang et al., 2015). Interestingly, we observed that cyclic food availability for several cycles does not result in consistent hyperactivity across all cycles. Flies that experienced 12 h of starvation per day for 7 consecutive days, showed activity levels comparable to the controls in the 12 h starvation window each day. However, activity levels on the first two days were higher as compared to other days (Supp Fig 3A, repeated measures ANOVA, Greenhouse-Geisser ε = 0.44, F_(2.67, 149.3)_ = 28.39, main effect of day, *p* < 0.001, followed by pair-wise *t*-tests with Bonferroni corrections for 21 comparisons). When flies were subjected to FS10:14, they showed an immediate increase in activity in response to lack of food (Fig 1B right, Supp Fig 3B). The activity levels were higher than the controls in the first 3 cycles of the treatment, after which they were comparable to the controls (Supp Fig 3B, Greenhouse-Geisser ε = 0.52, F_(3.13, 175.32)_ = 3.72, day × treatment, *p* = 0.012, followed by pair-wise *t*-tests with Bonferroni corrections for 49 comparisons). Interestingly, we found two types of behaviours among individuals when subjected to 16 h of starvation, 33.3% flies (7/23) showed excessive activity during starvation hours throughout the 8 days of FS 8:16 (type *a* flies, Fig 1C; centre) and 60.9 % flies, (14/23) appeared to show excessive activity during starvation only for the first few days after which the activity seemed to decrease (type *b* flies, Fig 1C; right). Day-to-day activity levels of FS8:16 flies (Supp Fig 3C, Greenhouse-Geisser ε = 0.29, F_(2.05, 108.54)_ = 6.99, day × treatment, *p* = 0.001, followed by pair-wise *t*-tests with Bonferroni corrections for 64 comparisons) showed reduction in activity levels after 6 days (Fig 1C right, Supp 3C). This shows that phases during FS are masked as a result of hyperactivity occurring in response to starvation.

### T26 and T21 Feeding: Starvation cycles do not synchronize the activity/rest rhythm of Drosophila melanogaster

We then assessed the third criterion for entrainment, i.e., period matching between activity/rest rhythm (τ) and external FS cycles (T). In the previous experiments given that T was 24 h, and τ was also close to 24 h (Table 1) it was not possible to test this criterion using data from FS12:12, 10:14 and 8:16 experiments. We therefore imposed FS cycles whose periods were not 24 h, but also not very deviant from 24 h to account for a possibility of a food entrainable oscillator if present, having narrow limits of entrainment. We imposed either a 26 h or a 21 h FS regime and asked if this could result in period matching of the activity/rest rhythm to external FS cycles. We subjected the flies to T26 FS cycles wherein the flies were provided food for 13 hours and were starved for 13 hours (Fig 5A) in an otherwise aperiodic environment (DD, 25°C). A large majority (Fig 5B) of the experimental flies appear to free run with a phase similar to the previous entrained phase in LD12:12 despite being subjected to T26 FS regime. However, they also exhibited a masked component to the disturbance caused due to change of tubes (Fig 5A, green dashed line). A Chi square periodogram analysis during T26 FS revealed 2 significant periodicities; one that was close to 24 h (henceforth referred to as free running component) and another that was close to 26 h (Table 2, Fig 5B). Since both control and experimental flies exhibit this long period (∼26 h) component, the same can be attributed to the physical disturbance experienced by both the controls and experimental flies (henceforth referred to as the masking component). Presence of two significant periodicities in a large proportion of experimental flies similar to control flies suggests that a T26 FS regime is unable to synchronise activity (Fig 5B, Chi-square test for proportions shows no difference in the two groups (*p* = 0.18, Chi-square = 3.22, *df* = 2). We estimated the difference in free running component (∼24h period) during FS (^24^ _τ1,_T26 FS) and after FS (^24^_τ2,_ D *ad lib*) for each fly. Interestingly, we found that experimental flies exhibited approximately 23 minutes longer period during T26 FS regime. This difference was significantly greater than the controls suggesting that T26 FS indeed significantly lengthens the internal period of the flies, albeit to a small degree (Fig 5C, U = 2388, *p* < 0.001, Mann-Whitney U test, Table 2).

**Figure 5:**
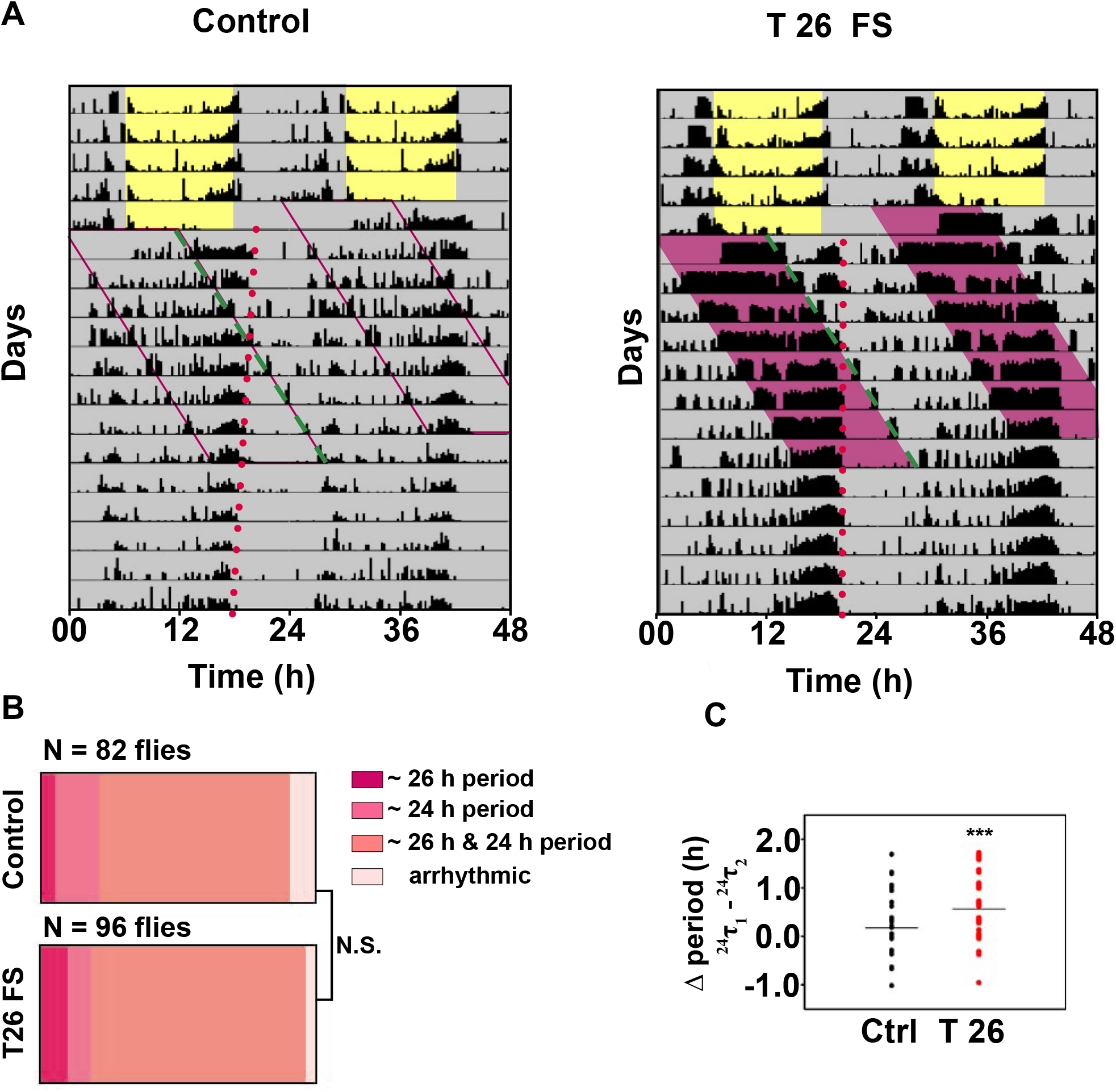
FS T26 cycles do not entrain the locomotor activity/rest rhythm. **(A)** Representative actograms of **(left)** control and **(right)** experimental flies recorded under LD12:12 conditions for the first 4 days before being subjected to T26 FS cycles in DD conditions. Pink shaded region represents 13 hours of starvation on each day. T26 FS cycles continued for 7 days followed by DD *ad libitum* conditions. Both the control and the experimental flies show masked responses to the T26 regime along with a free running component of activity (^24^ _τ1_) that seems to follow the phase from the LD12:12 cycle. Dashed line tracks the startle activity due to food changes (green) and dotted line tracks activity offset phase after LD 12:12 (red). **(B)** Proportion of flies showing two different periodicities (26 h and 24 h), only 26 h or only 24 h periodicity or arrhythmicity during T26 FS are indicated. A major proportion of both control and experimental flies show dual periodicities. **(C)** Distribution of changes in the ∼ 24 h periodicity during the T26 regime for controls and experimental flies. Differences were calculated by subtracting periods after T26 FS **(^24^_τ2_)** from the periods during T26 FS **(^24τ1^)**. T26 FS flies have a longer free-running component (∼24 h) during T26. NS denotes no difference between controls and T26 FS flies. *** denotes *p* < 0.001.

**Table:2.** Number of flies showing two periodicities when subjected to FS T26.

|  | n | No. of flies with 2 periodicities ~24h & ~26 | Flies exhibiting only ~ 26 h periodicity | Flies exhibiting only ~ 24 h periodicity | Flies exhibiting >2 periodicities | Arrhythmic flies | Average period (~24 h) | Average period (~26 h) |
| --- | --- | --- | --- | --- | --- | --- | --- | --- |
| Control | 94 | 57 | 5 | 13 | 7 | 7 | 23.94 ± 0.06 | 25.96 ± 0.02 |
| T26 | 120 | 75 | 10 | 8 | 14 | 3 | 24.27 ± 0.14 | 25.98 ± 0.004 |

Like the T26 regime, we also subjected the flies to a T21 regime with 10.5 h of food availability followed by 10.5 h of starvation in DD 25°C. We found similar results wherein the experimental flies continued to free run from the previously entrained LD phase along with a masked response to the external food availability cycles (Fig 6A, green dashed line). Akin to T26, we found flies exhibiting 2 periodicities, a near 24 h component (free running component) and a near 21 h component (masked component) as estimated by Chi square periodogram (Fig 6B). A higher proportion of T21 FS flies show only 21 h periodicity as compared to controls, (Chi-square test for proportions shows a significant difference *p* < 0.1, Chi-square = 15.37, *df* = 2) yet majority of the experimental flies still show both 21h and 24h periodicities. This further provided evidence for inability of the external T21 FS cycle to synchronize the activity/rest rhythm because of incomplete period matching. Moreover, like the T26 regime, T21 regime also affects the intrinsic period during the treatment. We estimated the difference in free running period during FS (^24^_τ1,_ T21 FS) and after FS (^24^_τ2,_ DD *ad lib*) for each fly and we found that experimental flies exhibited approximately 24 minutes shorter period during T21 FS regime. This difference was significantly lower than the controls suggesting that T21 FS significantly shortens the intrinsic period of the experimental flies (Fig 6C, U = 2389, *p* < 0.001, Mann-Whitney U test, Table 3). Overall, inability to match the intrinsic period to T cycles suggests that FS cycles are inefficient in synchronizing the activity/rest rhythm. Thus, another criterion for entrainment (period matching: =T) is not met under conditions where food availability is cyclic. These results taken together, suggest that FS cycles do not entrain the activity/rest rhythms controlled by the central clock in *D. melanogaster*.

**Figure 6:**
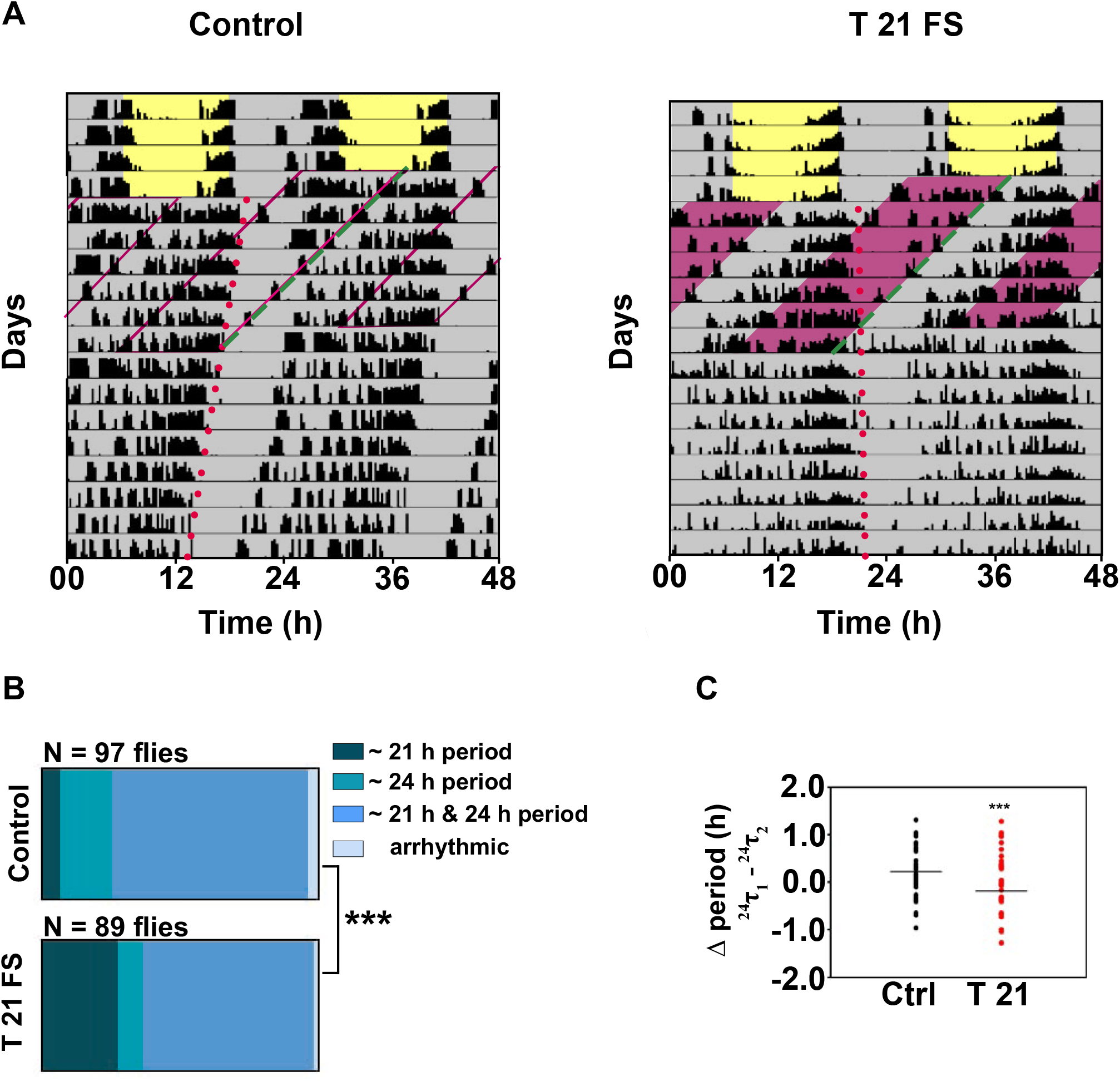
T21 cycles do not entrain the locomotor activity/rest rhythm in *D. melanogaster*. **(A)** Representative actograms of **(left)** control and **(right)** experimental flies recorded in LD 12:12 conditions for the first 4 days before being subjected to T21 FS cycles in DD. Pink shaded region represents 10.5 hours of starvation on each day. T21 FS cycles continued for 7 days followed by DD *ad libitum* conditions. Both the control and the experimental flies show masked responses to the T21 regime along with a free running component of activity (^24^_τ1_) that seems to follow the phase from the LD12:12 cycle. **(B)** Proportion of flies showing two different periodicities (21 h and 24 h), only 21 h or only 24 h periodicity or arrhythmicity during T21 FS are indicated. A major proportion of both control and experimental flies show dual periodicities. A higher proportion of T21 FS flies show only 21 h periodicity. **(C)** Distribution of changes in the ∼ 24 h period during the T21 regime for **(left)** controls and **(right)** experimental flies. Differences were calculated by subtracting periods after T21 FS **(^24^**_τ_**_2_)** from the periods during T21 FS **(^24^**_τ_**_1_).** T21 FS flies have a shorter free-running (∼24 h) component during T21. *** denotes p < 0.001. All other details as in Figure 5.

**Table:3.** Number of flies showing two periodicities when subjected to FS T21.

|  | n | No. of flies with 2 periodicities ~24h & ~21h | Flies exhibiting only ~ 21 h periodicity | Flies exhibiting only ~ 24 h periodicity | Flies exhibiting >2 periodicities | Arrhythmic flies | Average period (~ 24 h) | Average period (~ 21 h) |
| --- | --- | --- | --- | --- | --- | --- | --- | --- |
| Controls | 108 | 69 | 7 | 18 | 11 | 3 | 24.03 ± 0.05 | 20.97 ± 0.009 |
| T21 | 95 | 55 | 25 | 8 | 6 | 1 | 23.82 ± 0.03 | 21.01 ± 0.02 |

## DISCUSSION

We examined if cyclic food availability can act as an entraining cue for the clock in fruit fly *Drosophila melanogaster*. We assessed three criteria for entrainment - day-to-day phase stability, phase control and period matching. We found that subjecting flies to FS cycles of different durations of starvation does not result in inter-individual synchrony and phase control. These results build on the findings of a previous study in which subjecting flies to FS12:12 cycle with feeding restricted to night time did not affect activity/rest pattern (Oishi et al., 2004). However, in each of the T24 cycles, we found that day 1 of the treatment had a dramatic effect on phase which persisted for 3-4 cycles in FS10:14 and FS8:16. Furthermore, T26 and T21 FS cycles could bring about a significant lengthening and shortening of period while only partially synchronizing the activity/rest rhythm to the external periodicity. These results suggest that while food cannot act as an entraining agent, it affects the clock properties such as phase and period. This indicates that the clock is perceptive food availability in the environment and can respond by making small changes to clock properties without changing the overall pattern of activity/rest.

*Drosophila* larvae are known to feed voraciously until they acquire critical weight before pupation; as adults however, feeding is meager. Nevertheless, when deprived of food, flies are known to respond by increasing their locomotor activity (Knoppien et al., 2000); (Keene et al., 2010) which has been attributed to foraging behaviour in flies (Yang et al., 2015). Indeed, in all our experiments we observed SIH in both male and female flies. Since hyperactivity is a direct and immediate response to starvation, it can be considered as masking. In FS10:14 and FS8:16 regimes we found that SIH decreased over subsequent cycles. This masking to lack of food is also reflected in the phase changes observed during FS cycles. We observed that day-to-day phase changes indeed mirror the activity levels that were also higher than baseline during the first few cycles and tapered to baseline in the subsequent cycles. Since acrophase is a phase marker that depicts the radial centre of mass of activity, SIH may influence the day-to-day phases. This influence of cyclic starvation disappeared after release into constant conditions with *ad lib* food. This was evident from the fact that the Δ phase values were found to be significantly different from zero in all three T24 FS paradigms tested. Therefore, while food availability is unable to entrain the activity/rest rhythm, masking to starvation may bring about some changes which indirectly affects the phase of the clock. This is of significance to organisms that often encounter unpredictability in food availability in their environment. Masking to changes in food availability becomes more relevant for female flies who in addition to their own survival, also need food patches for laying eggs. In fact, when female flies were subjected to FS8:16 regime, most flies show excessive activity throughout the period of starvation (Supp fig 4) in contrast with male flies that displayed reduction in SIH midway into the treatment (Fig 1C, c’). Average acrophase during the FS treatment changes in female flies but they immediately revert to a value similar to pre-FS cycle when shifted to constant conditions. However, despite SIH and its influence on the acrophases, the activity/ rest rhythm continues to free run during the FS cycles. This suggests that in females like in males, masking and free-running components together regulate day-to-day phases.

If flies were consistently masking to lack of food by increasing their activity levels, we would expect a higher synchrony in inter-individual phases. However, we found that the inter-individual synchrony was lacking both in males as well as female flies irrespective of short or prolonged hyperactivity response. Moreover, this increased activity was not only variable across days but also variable across individuals. Therefore, unlike typically masked responses which consists of consistent all-or-none responses, in our paradigm we observed a graded response to lack of food. This graded response may in part explain why inter-individual synchrony is absent despite occurrence of masking.

Food availability has been shown to affect activity patterns of many insects. Solitary bee species such as carpenter bees and orchid bees show foraging behaviour which is partially regulated by circadian clocks (Bloch et al., 2017). Social honey bees *Apis mellifera* entrain to food availability in their environment (Frisch and Aschoff, 1987) underlining the importance of circadian clocks in anticipating availability of food. Blood mealtimes are carefully phased to host availability in various hematophagous species such as bed bugs, kissing bugs and mosquitoes. For example, bed bug *Cimex lectularius* are active during dawn, a time when the humidity is relatively high and hosts are resting (Barrozo et al., 2004). Similarly, kissing bug *Triatoma infestans*, the vector for Chagas disease shows peak locomotion, feeding, and carbon-dioxide sensitivity in the early night presumably when hosts are inactive (Barrozo et al., 2004). Flight activity and feeding/biting patterns in mosquitoes are influenced by many environmental factors apart from the internal clock mechanisms (Barrozo et al., 2004). However, these feeding patterns have been shown to change in nocturnal *Anopheles spp* from peak feeding occurring during the nighttime to a relatively earlier phase due to change in host availability as a result of interventions to prevent spread of malaria in some African countries (Reddy et al., 2011); (Gatton et al., 2013);(Sougoufara et al., 2017). Such flexible patterns are also observed in *Culex pipiens* when blood meals were restricted to a daytime window (Fritz et al., 2014). Ground beetle *Feronia madida*, have also been previously shown to change their nocturnal activity patterns to diurnal activity patterns under starved conditions (Williams, 1959); (Beck, 1980). Cockroach *Periplaneta americana* that have nocturnal activity and feeding show masking response to food availability when food is restricted to daytime, however, the activity is still higher during the night suggesting little change to overall activity/rest pattern (Beck, 1980). Therefore, non-alignment of food availability to an active state of the organism can influence the activity patterns and possibly the underlying clock. Additionally, mice gauge the food reward to predation risk ratio and accordingly change their temporal niche depending upon the environmental conditions (van der Vinne et al., 2019). Thus, activity patterns may change from nocturnal to diurnal in response to change in food availability and predation pressure. We sought to ask if similar pressure in the form of restricted food access to a certain time of day can bring about a change in activity/rest of *Drosophila melanogaster*. We found that while the clock does not entrain to the food availability cycle it perceives these changes in food availability and accommodates them by adjusting its clock properties. This means that even though the activity may free run during the FS cycles, it free runs with a different phase. Different extent of clock responses to food could be because of difference in amount of food required for sustenance and/or difference in natural history and ecology of these species.

In mammals, anticipatory response to food availability suggests that circadian clocks are directly responsive to food availability. While the identity of such a clock (FEO) remains elusive to date, it is clear that FEO controls food anticipatory activity (Mistlberger, 1994); (Stephan, 2002). In *Drosophila*, response to restricted food access is masked, which suggests a homeostatic control. Our study demonstrates that this masked response in the form of SIH, indirectly affects the activity-controlling central clock. The physiological basis for this interaction between the homeostatic components and clock components nevertheless has not been studied so far. Future studies directed at understanding this interaction between clocks and a food homeostat will help to understand how food availability in the environment can shape the activity/rest patterns in animals.

## Acknowledgements

We thank Srikant Venkitachalam for help with statistics, Chitrang Dani for comments on the manuscript and other lab members for discussions. We also thank Rajanna Narasimhaiah and Muniraju Muniappa for technical assistance. This work was supported by intramural funds of Jawaharlal Nehru Centre for Advanced Scientific Research and and consumable grant from the Department of Biotechnology (DBT), Government of India, to V.S. (BT/ INF/22/SP27679/2018).

## Data availability statement

**The data that support the findings of this study are available from the corresponding author upon reasonable request.**

## Supplementary online material

**Supplementary Figure 1:**
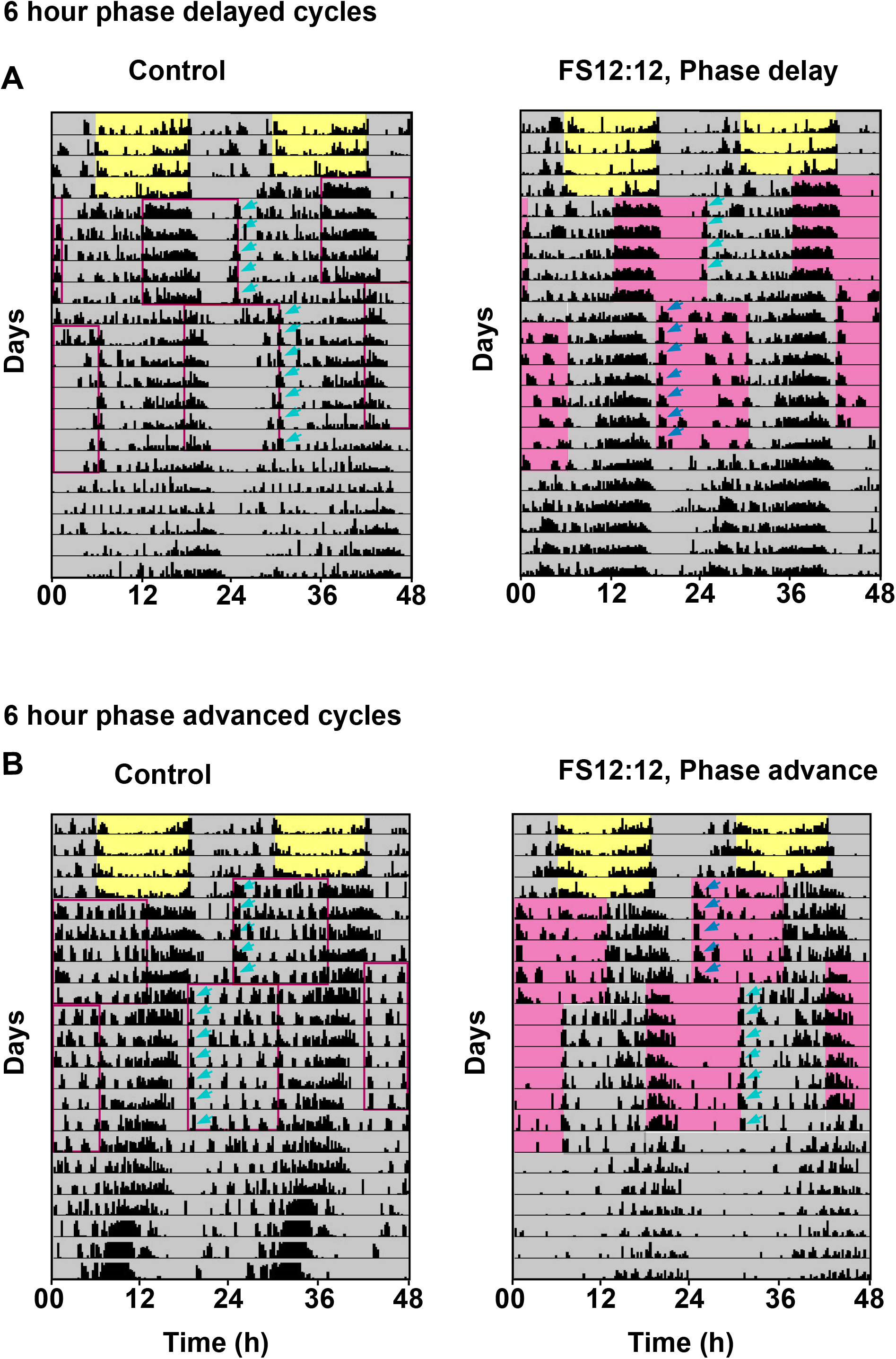
Phase delayed and phase advanced FS cycles do not synchronize the locomotor activity/rest rhythm in *D. melanogaster*. **(A)** Representative double plotted actograms of age matched disturbance control flies **(left)** and flies subjected to two consecutiveFS12:12 cycles phase delayed to one another **(right)**. Flies were housed in LD12:12 and later subjected to FS12:12 (pink shaded region depicts starvation hours) for 5 cycles. Starvation period **(FS1)** began 6 hours phase delayed with respect to the LD12:12 such that the LD and the FS regime were not in-sync with each other. FS1 cycles were further delayed by another 6 hours after the first 5 days, and this continued for another 7 days **(FS2)**. Flies were subsequently released into constant conditions with *ad libitum* food (DD, 25°C). Arrows indicate the startle bouts. **(B)** Representative double plotted actograms of age matched disturbance control flies **(left)** and flies subjected to two consecutiveFS12:12 cycles phase advanced to one another **(right)**. Flies were subjected to FS12:12 for 5 cycles in DD. **FS1** was 6 hours phase advanced compared to the initial LD12:12 regime. FS1 was followed by **FS2** which was further advanced by 6 hours compared to FS1. All other details are same as **(A)**. Experimental flies fail to synchronise to FS1 and FS2 in both phase delay and phase advance experiments and continue to display a phase from the previous LD cycle.

**Supplementary Figure 2:**
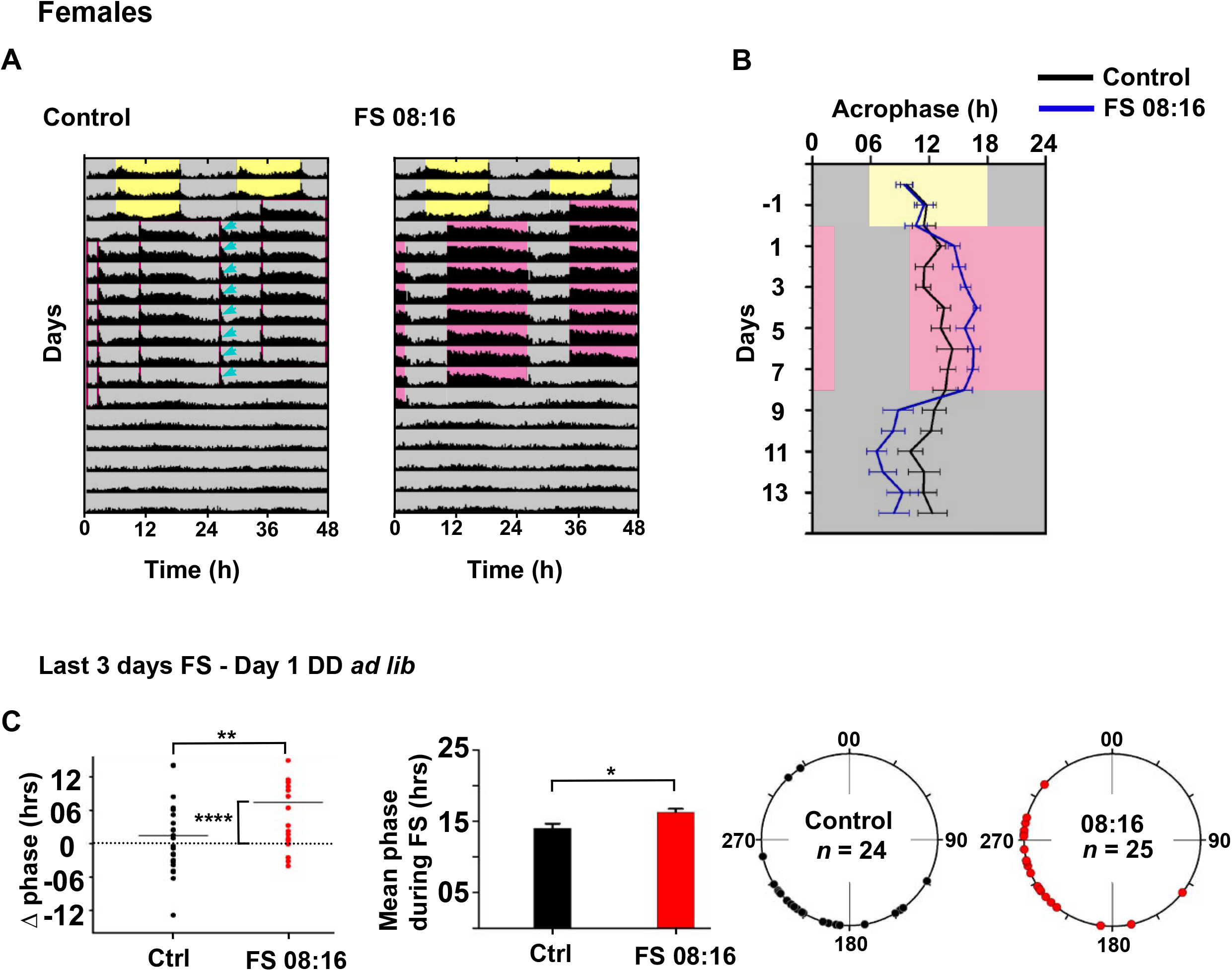
Female flies do not get entrained to FS8:16 cycles. **(A)** Double plotted batch actograms of **(left)** age matched disturbance control flies and **(right)** female flies subjected to FS8:16 regime. Flies show a prolonged activity bout on all cycles of the treatment. **(B)** Mean acrophases ± SEM of controls (black line) and FS flies blue line) across days of LD 12:12, FS8:16 treatment and DD *ad lib*. Experimental flies show phase change during the regime. Phases revert to previous values immediately after DD *ad lib*. **(C) (Left)** scatter plots showing Δ phases of individual control (black) and experimental (red) flies, *, ** indicate *p* < 0.05, 0.01 respectively. **(Centre)** Bar graph depicts mean acrophase (h) ± SEM during FS8:16 of disturbance controls (black) and experimental (red) flies in the last 3 days. (**Right)** polar plots depicting average acrophases for the last 3 days of individual flies during FS 8:16. **(left)** black circles indicate phases of control flies and **(right)** red circles indicate the phase of experimental female flies that were subjected to FS8:16 regime. All other details as in Figure 1 & 3.

**Supplementary Figure 3:**
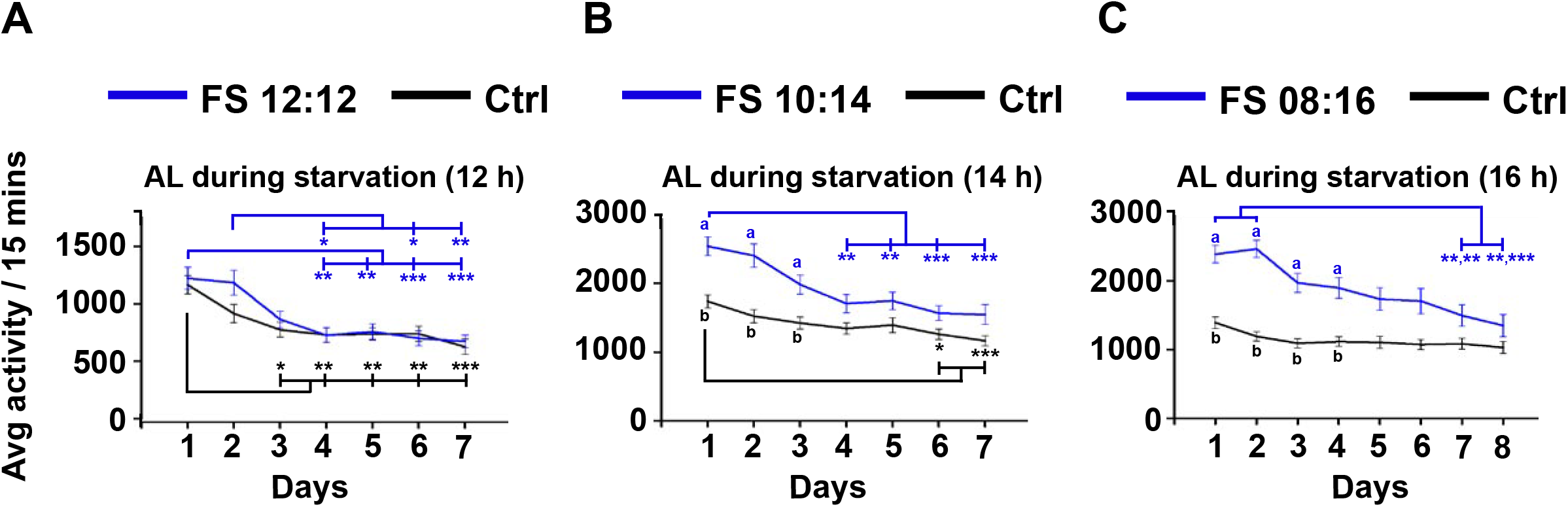
FS12:12, 10:14, 08:16 cycles do not result in sustained starvation-induced hyperactivity throughout the period of FS regime. Activity levels during starvation period of each day ± SEM during **(A)** FS12:12, **(B)** FS10:14, **(C)** FS8:16. Starvation induced hyper-activity in the experimental flies progressively reduces during the latter half of the 10:14, 8:16 FS cycles. Letters denote significant differences (p <0.05) between controls and FS flies. *, **, *** denote *p* < 0.05, 0.01, 0.001 respectively.

